# NRF2-Dependent Anti-Inflammatory Activity of Indole via Cell Surface Receptor Signaling in Murine Macrophages

**DOI:** 10.64898/2025.12.21.695859

**Authors:** Clint Cheng, Siddanagouda R. Shivanagoudra, Aniruddh Mukunth, Sumathi Biradar, Yadeliz Serrano Matos, Cory Klemashevich, Da Mi Kim, Gus A Wright, Sweta Ghosh, Venkatakrishna Rao Jala, Stephen H. Safe, Robert C. Alaniz, Arul Jayaraman

**Affiliations:** Artie McFerrin Department of Chemical Engineering, Texas A&M University, College Station, TX; Department of Biomedical Engineering, Texas A&M University, College Station, TX; Veterinary Medicine & Biomedical Sciences, Texas A&M University, College Station, TX; Department of Microbiology and Immunology, University of Louisville School of Medicine, Louisville, KY; Department of Veterinary Physiology and Pharmacology, Texas A&M University, College Station, TX; Department of Microbial Pathogenesis and Immunology, College of Medicine, Texas A&M University, College Station, TX

## Abstract

In this study, we report indole’s anti-inflammatory effects to be AhR-independent in RAW 264.7 macrophages. To explore the possibility of indole’s surface-receptor mediated signaling, we developed an indole-bovine serum albumin conjugate (I3B), which primarily engage cell surface receptors and has limited intracellular engagement. Treatment with 10 μM of I3B led to a comparable reduction of TNF-α production in LPS-stimulated RAW 264.7 macrophages to that observed with 500 μM of free indole. Transcriptome profiling of I3B-treated LPS-stimulated RAW 264.7 macrophages revealed, I3B blunts pro-inflammatory response and induces gene signatures consistent with NRF2 activation. LPS-stimulated NRF2^-/-^ Bone Marrow-derived Macrophages (BMM) treated with I3B, showed higher levels of pro-inflammatory cytokine production relative to non-treated BMM. To define the upstream pathways responsible for this NRF2-depedent response, we examined GPCR-mediated signaling and found that I3B engages a Gq-coupled receptor to induce PKCδ phosphorylation, and subsequent NRF2 phosphorylation. Our results suggest I3B signals through a surface-receptor in a NRF2-dependent manner to reduce inflammation in murine macrophages.

**Teaser:** A cell-impermeant indole conjugate inhibits inflammatory signaling in macrophages through an NRF2-dependent mechanism.

## Introduction

Indole is a tryptophan-derived metabolite produced by intestinal microbes harboring the enzyme tryptophanase, TnaA (*1*). Metagenomic analysis of human gastrointestinal (GI) microbes suggest numerous phyla of bacteria, including *Firmicutes*, *Bacteroidetes*, and *Proteobacteria*, have genes encoding TnaA and can potentially produce indole (*2*). In human fecal samples, the median indole concentrations is approximately in the low millimolar range, with one study reporting values ranging from 0.3 to 6.64 mM in healthy adults (*3, 4*). Several *in vitro* and in *vivo* studies have demonstrated a role for indole in modulating inflammation across multiple models. These include suppressing diet-induced NAFLD and liver inflammation (*5, 6*), protecting against dextran sodium sulfate-induced colitis and promoting intestinal epithelial function (*7, 8*), and reducing central nervous system inflammation in a murine model of multiple sclerosis (*9*). In addition to indole, other microbially derived tryptophan metabolites (MDTMs), including indole-3-aldehyde (IAld), indoxyl-3-sulfate, indole-3-propionic acid, indole-3-acetate, indole-3-ethanol, and indole-3-pyruvate, have also demonstrated similar protective effects in various model systems (*10*).

Several studies have shown that indole and other MDTMs signal through the cytosolic aryl hydrocarbon receptor (AHR) pathway in different cell types, such as epithelial cells and hepatocytes (*10, 11*). However, AhR-dependent effects can vary across cell type, and it remains unclear whether MDTMs engage other receptors with different levels of activity (*12*). Although studies have shown MDTMs modulate macrophage signaling (*6, 10*), the AhR-dependent activity of these compounds is yet to be fully elucidated. Certain MDTMs, notably kynurenic acid and tryptamine, have been shown to activate G-protein coupled receptors (GPCRs), such as GPR35 and 5-HT4R (*13, 14*), respectively. This suggest MDTMs can interact with cell surface receptors. However, evidence for a direct interaction between indole and cell surface receptors remains limited.

In this study, we investigated the effect of indole on inflammatory signaling in murine macrophages and characterized indole’s AhR dependence using an AhR antagonist and cells from AhR^-/-^ mice. We further explored the potential for indole to engage cell surface receptors, as opposed to cytosolic receptors like the AhR, by synthesizing a membrane impermeant indole-bovine serum albumin conjugate (I3B) and examined this compound ability to modulate LPS-mediated inflammatory signaling in RAW 264.7 macrophages. Lastly, we investigated the mechanism(s) underlying I3B signaling in RAW 264.7 macrophages.

## Results

### Indole modulates TNF-α production in RAW264.7 macrophages independent of the AhR

The effect of indole on LPS-stimulated TNF-α production in RAW 264.7 murine macrophage was assessed by intracellular cytokine staining (ICS) flow cytometry. RAW 264.7 macrophages pre-incubated with indole for 4 h, followed by 6 h of co-stimulation with 250 ng/mL of LPS, had significantly attenuated TNF-α accumulation relative to vehicle control in a dose-dependent manner, as determined by both Mean Fluorescence Intensity (MFI) and percentage of TNF-α positive live cells (**Fig. 1A-B**). At concentrations of 0.5 and 1 mM, indole reduced the proportion of TNF-α positive cells by roughly 26% and 43%, respectively (**Fig. 1B** and **C**). The decrease in TNF-α accumulation was also observed in primary bone marrow derived macrophages (BMM) from both WT and AhR^-/-^ mice, albeit with reduced potency (**Fig. 1D** and **E**). Incubation with 1 mM indole resulted in roughly 16% and 27% reduction in TNF-α positive cells in BMM from WT and AhR^-/-^ mice, respectively. No statistically significant differences were observed between the WT and AhR^-/-^ conditions, suggesting indole inhibits LPS-stimulated TNF-α expression in macrophages in an AHR-independent manner.

**Fig. 1.**
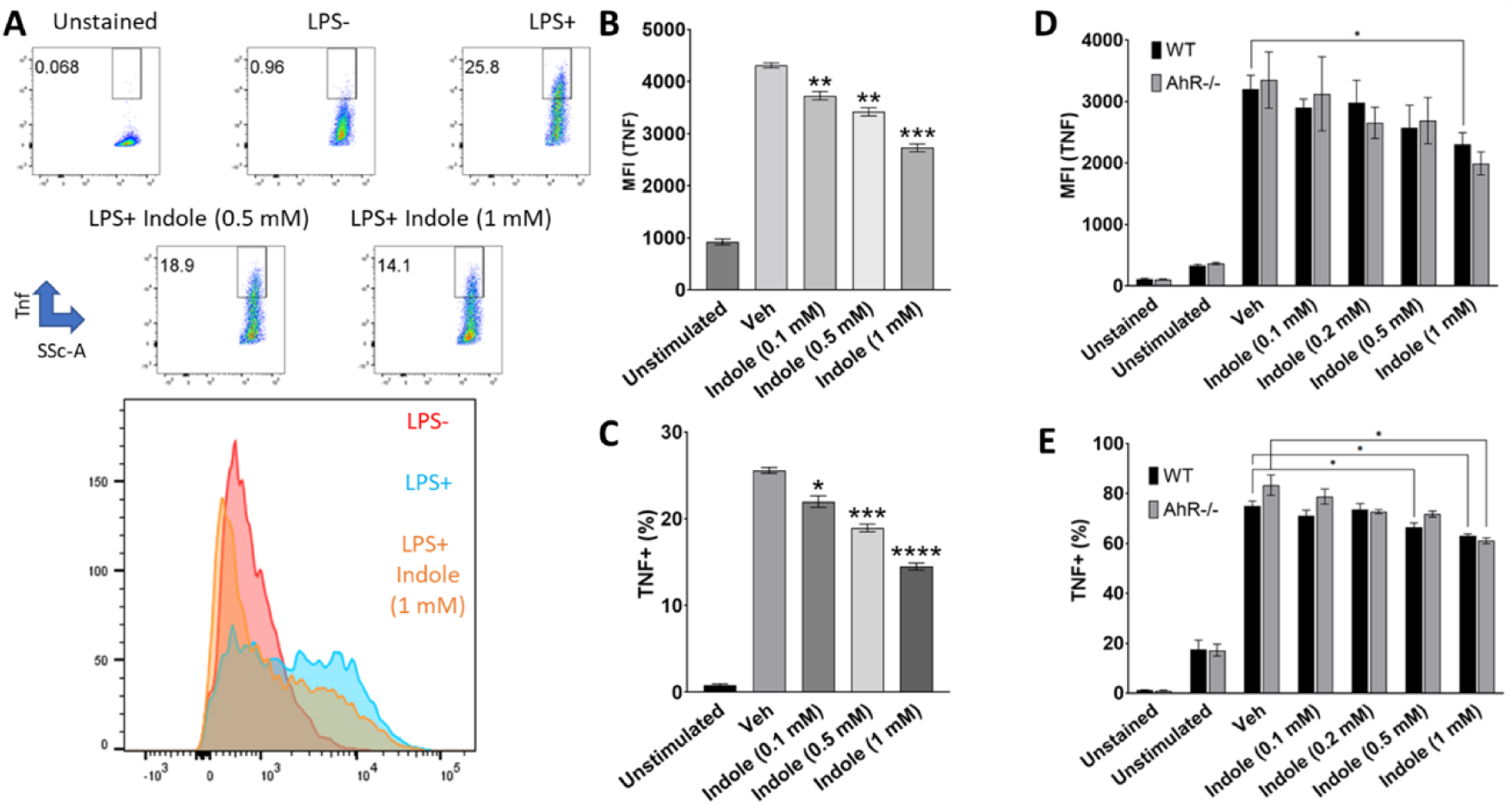
Indole inhibits LPS stimulated TNF accumulation independent of the AhR. RAW264.7 macrophages were pre-incubated with indole or vehicle control for 4 h prior to co-stimulation with 250 ng/mL of LPS for 6 h. **(A)** Representative gating strategy, scatter plots, and histograms of RAW264.7 cells analyzed with ICS flow cytometry. **(B)** TNF-α protein Mean Fluorescence Intensity (Arbitrary Units) and **(C)** Percent TNF-α positive cells from RAW 264.7 cells. **(D)** BMM from WT or AhR^-/-^ mice were pre-incubated with indole or vehicle control for 4 h prior to co-stimulation with 20 ng/mL LPS for 6 h. Cells were stained for TNF-α protein and analyzed using ICS flow cytometry and quantified using the gating strategy shown in **(A)**. TNF-α protein mean fluorescence intensity (arbitrary units) and **(E)** percent TNF-α positive cells. Bars represent mean values from three or four independent cultures and error bars represent ± S.E.M. * *p* < 0.05, ** *p* < 0.01, *** *p* < 0.001, **** *p* < 0.0001.

### Cell-impermeant indole potentiates the anti-inflammatory properties of free indole

Based on the known interaction between MDTMs such as kynurenic acid and serotonin with cell surface receptors and indole’s AhR-independent modulation of TNF-α expression in murine macrophages, we hypothesized that indole engages cell surface receptors to attenuate LPS-mediated signaling. To test the cell surface localized effects of indole, we conjugated indole-3-aldehyde (IAld) to bovine serum albumin (BSA) using the crosslinker, N-κ-maleimidoundecanoic acid hydrazide (KMUH), resulting in a 1:1:1 stoichiometry between an indole moiety, KMUH, and BSA (**Fig. 2A**). The end-product, indole-3-BSA (I3B), comprises of an indole moiety linked to one end of an 11-carbon aliphatic spacer via a hydrazone bond, with a thioether bond linking BSA at the opposing end of the spacer. The predicted mass shift of I3B relative to BSA is 423 Da, and MALDI-TOF mass spectrometry analysis showed an observed mass shift of 449 Da (**Fig. 2B**).

**Fig. 2.**
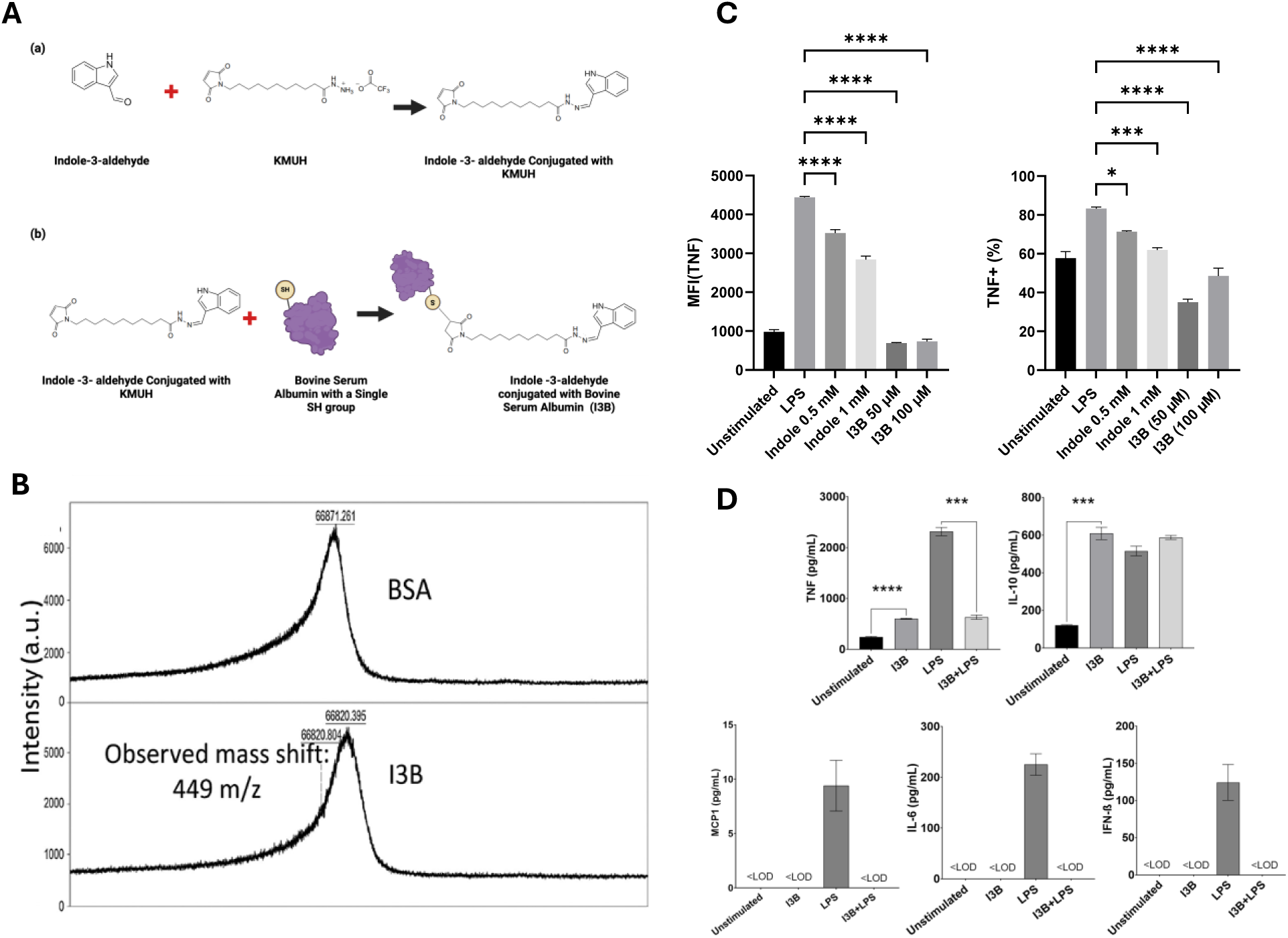
Synthesis and Characterization of I3B and Its Anti-inflammatory Effects. **(A)** Indole-3-aldehyde (IAld) was reacted with KMUH in DMSO at a 5:1 molar ratio. The reaction product was further reacted with BSA at a 5:1 molar ratio to obtain I3B. **(B)** MALDI-TOF mass spectrometry characterization of desalted I3B and BSA. **(C)** RAW264.7 macrophages were pre-incubated with compounds of interest for 4 h followed by co-stimulation with 250 ng/mL of LPS for 6 h and ICS flow cytometry. Quantification of TNF-α protein was carried out using the gating strategy shown in Figure 1(A). Mean fluorescence intensity (arbitrary units) and percent TNF-α positive cells are shown. (**D**) Supernatants from macrophages pre-incubated with 100 µM I3B for 4 h, then co-stimulated with 250 ng/mL of LPS for 24 h, were analyzed with a LegendPlex^TM^ multiplexed antibody bead-based assay. Data are from three independent cultures with two technical replicates for each culture. Error bars are ± S.E.M. * *p* < 0.05, ** *p* < 0.01, *** *p* < 0.001, **** *p* < 0.0001, <LOD: below limit of detection.

Pre-incubation with 10 μM of I3B attenuated the LPS stimulated TNF-α accumulation in RAW264.7 macrophages to a level comparable to that observed with 1 mM free indole, while higher concentrations of 50 and 100 μM of I3B reduced TNF-α accumulation to levels lower than the unstimulated controls (**Fig. 2C**). Pre-incubation with free indole, IAld, or BSA (alone or in combination) at equivalent concentrations did not recapitulate the potent effects of I3B, suggesting this potentiation results from increased indole engagement with the cell surface (**Fig. S1**).

To further investigate the effects of I3B on the modulation of pro-inflammatory cytokine production, RAW 264.7 macrophages were pre-incubated with 100 μM of I3B for 4 h followed by LPS stimulation for 24 h. Cell culture supernatants were then analyzed via LEGENDPlex^TM^ using their Mouse Inflammation panel. Pre-incubation with I3B significantly reduced secretion of TNF-α, recapitulating the ICS flow cytometry results of free indole. Unstimulated RAW264.7 cells did not secrete detectable levels of IL-6, IFN-β, and MCP-1 compared to LPS stimulated cells which showed an increase in the concentration of these pro-inflammatory cytokines (225, 124, and 9 pg/mL, respectively after 24 h) (**Fig. 2D**). I3B pre-treatment on the other hand, led to a reduction in the concentration of these pro-inflammatory cytokines below the limit of detection. Intriguingly, incubation with I3B alone stimulated a five-fold increase in the secretion of anti-inflammatory cytokine IL-10 (from 122 pg/mL to 610 pg/mL). These results suggest I3B pre-incubation attenuates RAW 264.7 macrophage response to LPS consistent with an anti-inflammatory response. We hypothesized that the transport of a larger molecule such as I3B compared to a small hydrophobic molecule like indole (117 Da), will be reduced due to steric hindrance from the conjugated BSA (66 kDa). To demonstrate this, we synthesized a second molecule using similar chemistry as that used for I3B, except with used NeutrAvidin as the conjugate instead of BSA for fluorescence imaging. After 4 h of exposure, the indole-conjugated molecule was localized to the cell surface (**Fig. 3A** and **B**); supporting the hypothesis that indole conjugated to a larger protein (60 kDA) is excluded from RAW 264.7 macrophages. Fluorescence imaging further confirms that I3B’s anti-inflammatory effects are mediated through the engagement of cell surface receptors.

**Fig. 3.**
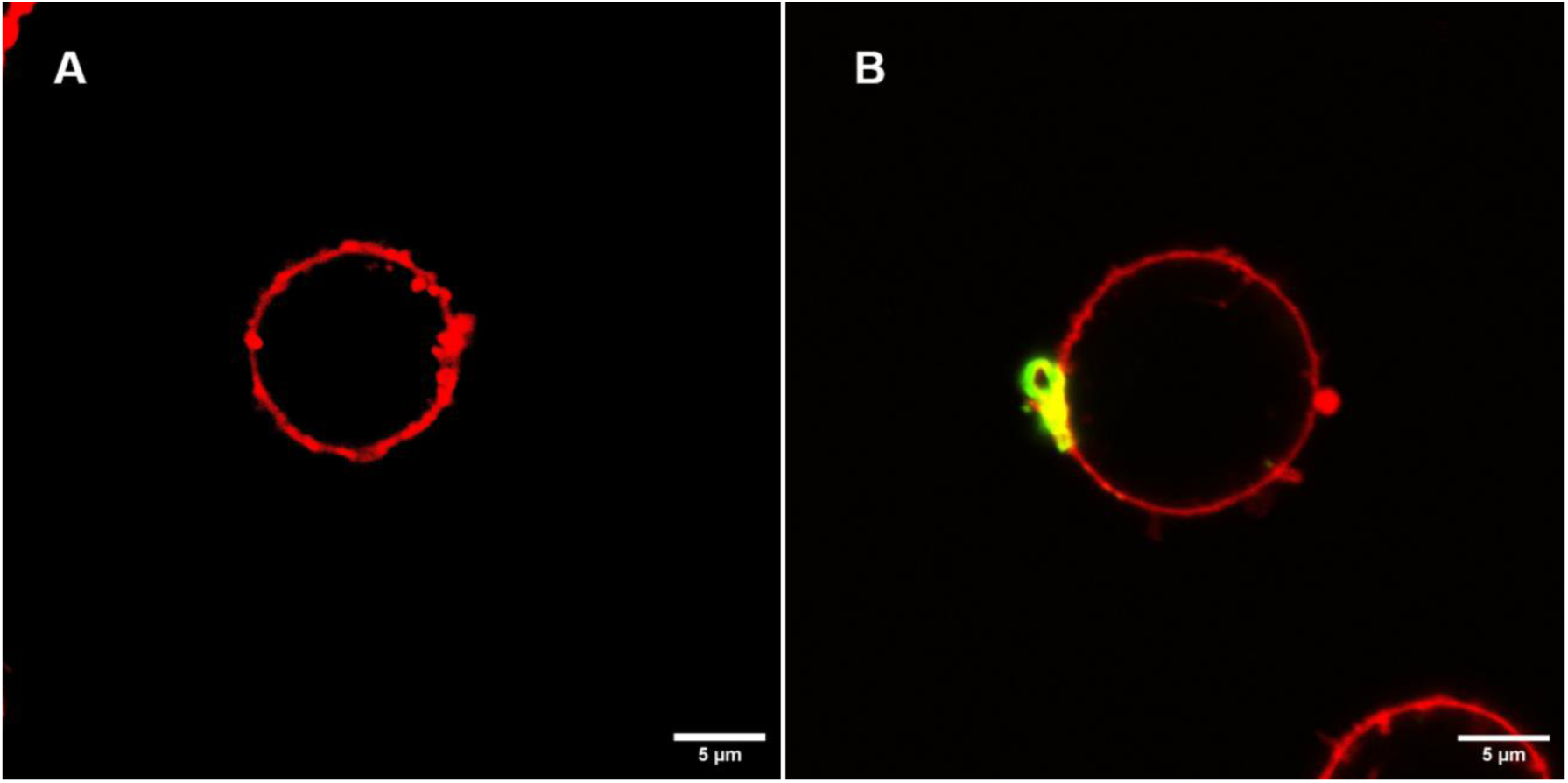
Cellular localization of Indole-NeutrAvidin conjugate following 4-hour incubation. Confocal microscopy images show that the Indole-NeutrAvidin conjugate (20 µM) remains confined to the cell surface after 4 hours, without evidence of intracellular uptake. **(A)** NeutrAvidin alone (20 µM) **(B)** Indole-NeutrAvidin conjugate (20 µM) demonstrates distinct membrane-associated localization.

### Effects of I3B in macrophages are independent of the AhR

We investigated the potential role of AhR in I3B signaling in macrophages. Bone marrow-derived macrophages (BMM) isolated from wild-type (WT) and AhR knockout (AhR^-/-^) mice were incubated with 50 or 100 µM of I3B, followed by co-stimulation with LPS. BMM treated with I3B, regardless of concentration, displayed a significant reduction in TNF-α compared to vehicle control (**Fig. 4A**). Importantly, no significant differences in TNF-α levels were observed between WT and AhR^-/-^ BMM. Furthermore, treatment of BMM with 50 or 100 µM I3B resulted in significantly greater inhibition of TNF-α compared to those seen with free indole, IAld, and BSA alone **(Fig. S2A)**. These results suggest that I3B inhibits LPS-stimulated TNF-α production via an AhR-independent mechanism in BMM. To test this, RAW264.7 cells were pre-incubated with 20 μM of the AhR inhibitor CH-223191 for 14 h, followed by co-incubation with I3B for 4 h and subsequent stimulation with 250 ng/mL of LPS for 6 h. While CH-223191 alone increased the MFI of TNF-α and the percentage of TNF-positive cells relative to vehicle control (**Fig. 4B**), the inhibitory effect of I3B on TNF-α production was comparable (79%) regardless of CH-223191 treatment. This AhR-independent inhibition mirrored observations with indole and IAld, though their magnitudes of inhibition were lower (19% and 42%, respectively, for indole and IAld) **(Fig. S2B).**

**Fig. 4.**
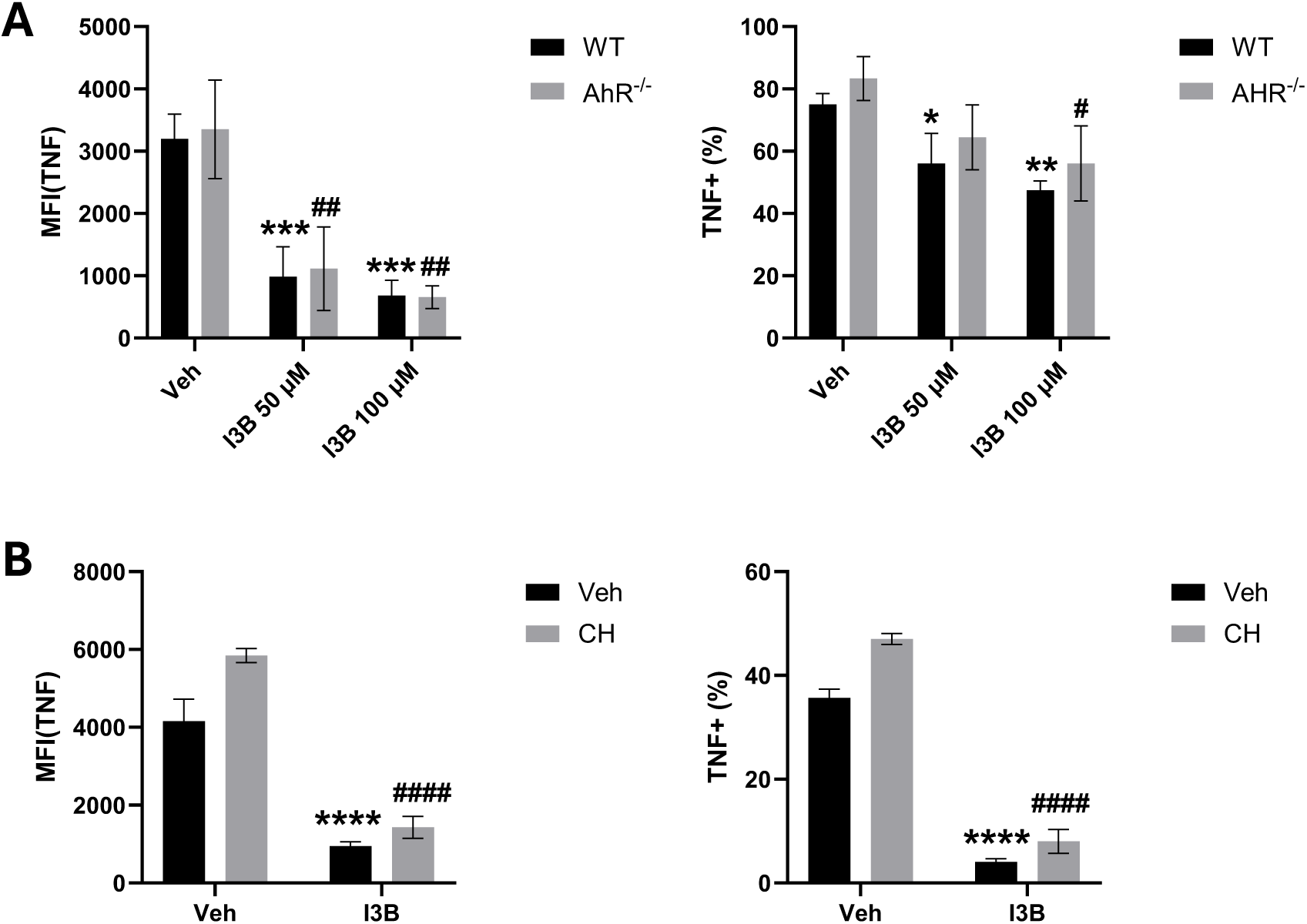
Effects of I3B on macrophages is AhR independent. (**A**) Bone marrow-derived macrophages (BMMs) from wild-type (WT) and AhR knockout (AhR⁻/⁻) mice were treated with I3B 50 and 100 µM or controls for 4 hours, followed by LPS stimulation (20 ng/mL) for 6 hours. TNF-α levels were assessed by ICS flow cytometry. Absolute MFI (left) and percent TNF-α–positive cells (right) are shown. (**B**) RAW264.7 macrophages were pre-treated with the AhR antagonist CH-223191 (CH; 20 µM) for 14 hours, followed by co-incubation with I3B at 50 µM, or vehicle control (Veh) for 4 hours. Cells were subsequently stimulated with LPS (250 ng/mL) for 6 hours and analyzed by intracellular cytokine staining (ICS) and flow cytometry. The absolute mean fluorescence intensity (MFI) values and percentages of TNF-α–positive cells. Data are presented as mean ± SEM. \**p* < 0.05, \*\**p* < 0.01, \*\*\**p* < 0.001 relative to vehicle control (A) or WT (B); #*p* < 0.05, ##*p* < 0.01, ###*p* < 0.001 relative to vehicle + CH condition (A) or AhR^-/-^ (B); ns = not significant.

### Transcriptional analysis of I3B-treated RAW macrophages

We performed RNA-seq in I3B pre-treated or not LPS-stimulated RAW 264.7 macrophages to elucidate how I3B regulates global gene expression. RAW 264.7 macrophages were incubated with or without 50 µM I3B for 4 h, followed by incubation with or without 250 ng/mL of LPS for 1 h. Multidimensional scaling of transcriptomic data revealed three distinct clusters, consisting of the unstimulated control, LPS-stimulated cells, and both I3B conditions (I3B with and without LPS) (**Fig. 5A**). As expected, LPS stimulation upregulated the gene expression of pro-inflammatory cytokines (**Table S1**). Among these, Cxcl2 and Cxcl10 exhibited the highest fold-changes (88.6 and 40.3-fold, respectively), followed by Egr2 (21.6-fold) and Tnf (10.1-fold). However, when cells were pre-treated with I3B, subsequent stimulation by LPS did not result in a significant induction in the gene expression of these cytokines (**Fig. 5B**).

**Fig. 5.**
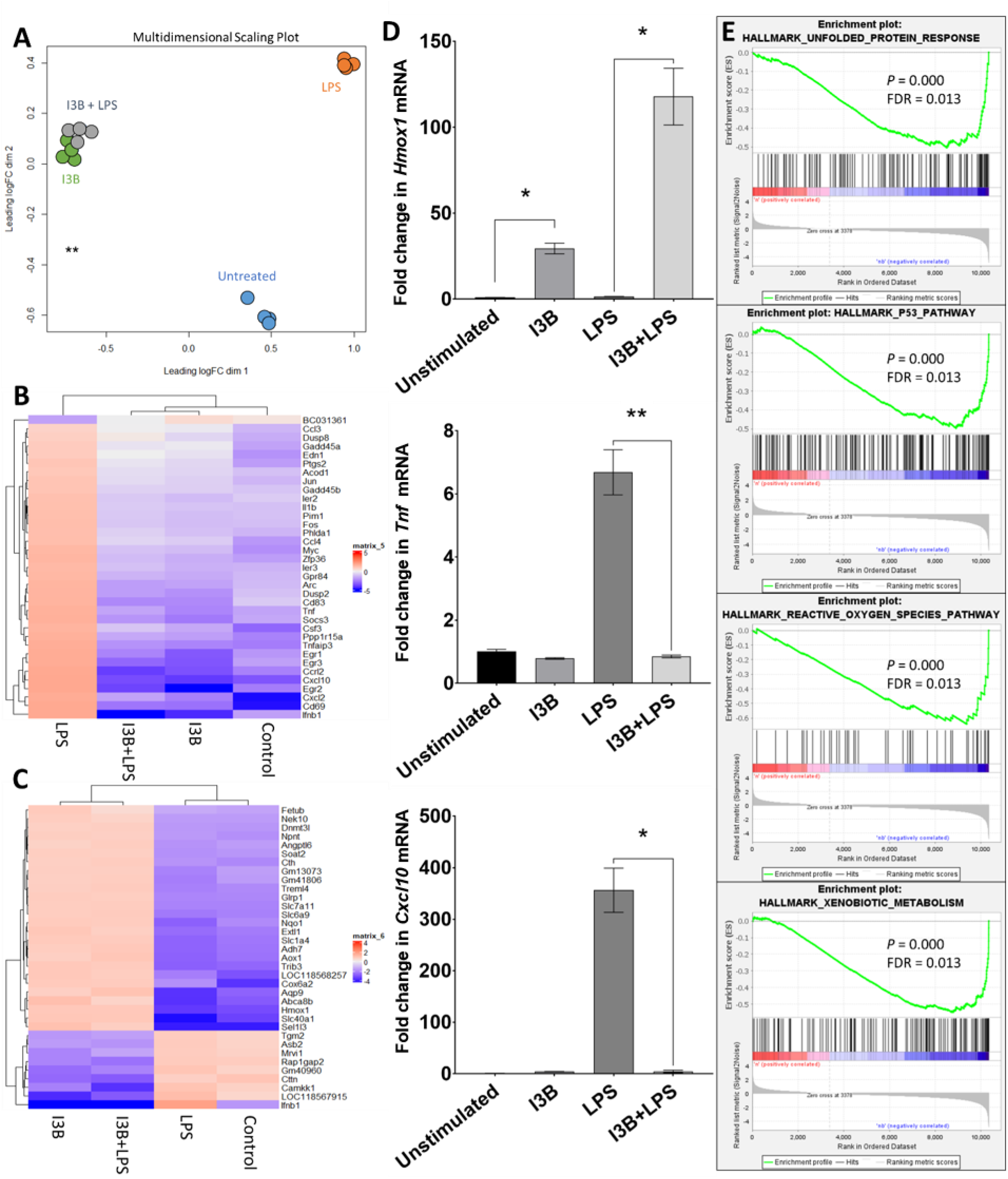
Transcriptome profiling of I3B treated RAW264.7 Macrophages. Cells were pre-incubated with 50 µM of I3B for 4 h followed by stimulation with 250 ng/mL LPS for 1 h. Transcript libraries were prepared for paired-end RNA-sequencing with differential gene expression analysis done in EdgeR. (**A**) Multidimensional scatterplot representing the four treatment conditions with four independent replicates. Clustered heat maps of the top 35 most differentially regulated genes by (**B**) LPS stimulation and by (**C**) I3B treatment alone. Differential gene expression data were row normalized and log2 transformed. (**D**) qRT-PCR validation of select differentially regulated genes (n = 3). (**E**) Gene set enrichment analysis barcode plots indicate the position of the genes in each gene set with blue colors representing those with positive Pearson correlations upon I3B treatment. Enrichment plots for the four hallmark gene sets with the highest normalized enrichment score are shown. * *p* < 0.05, ** *p* < 0.01.

I3B alone upregulated the expression of 334 genes and downregulated that of 77 genes (fold change ≥ 2 and *p* < 0.05) relative to the unstimulated control. This set of genes was distinct from that altered by LPS (**Table S2**). The list of genes up regulated by I3B included *Sel1l3*, *Cox6a2*, *Slc40a1*, and *Hmox1* (34.5, 23.2, 20.4, and 17.7-fold, respectively). Stimulation of I3B pre-conditioned cells with LPS largely did not alter the transcriptome of this set of genes (**Fig. 5C**). Patterns of gene expression in *Tnf*, *Hmox1*, and *Cxcl10* were validated using qRT-PCR (**Fig. 5D**).

### Gene set enrichment analysis

To further analyze the transcriptome data, Gene Set Enrichment Analysis (GSEA) was performed using two gene set collections: the Hallmark gene sets from the Molecular Signatures Database and the Wikipathway collection (*15*). GSEA showed that incubation with I3B enriched 18 of the 49 gene sets in the hallmark gene set collection (nominal *p* < 0.01, FDR < 0.2). Gene sets with the highest normalized enrichment score (NES) were the unfolded protein response (2.52), P53 pathway (2.50), reactive oxygen species pathway (2.38), and xenobiotic metabolism (2.35) (**Fig. 5E**). Analysis of the Wikipathway collection revealed enrichment in 71 of the 337 gene sets (nominal *p* < 0.01, FDR < 0.2) (**Table S3 and Fig. S3)**. Of particular interest was the NRF2-ARE (nuclear factor, erythroid 2-like 2-antioxidant response element) regulation gene set (NES = 2.48, rank 5), that contains multiple genes for phase II metabolizing enzymes upregulated by I3B, such as *Hmox1*, *Slc7a11*, *Nqo1*, and *Gclm* (18, 8, 7, and 5-fold upregulation, respectively).

### Anti-inflammatory effects of I3B are partially NRF2-dependent

The significant upregulation of Hmox1 transcript levels (17.7-fold), its presence in both the reactive oxygen species pathway and NRF2-ARE regulation gene sets, and previous research supporting Hmox1’s role in NRF2-regulated pathways (*16, 17*) prompted us to further investigate the contribution of NRF2 in I3B signaling. BMM were isolated and cultured from WT and NRF2^-/-^ mice and pre-incubated with 20 μM I3B prior to LPS stimulation. I3B pre-treatment reduced the LPS-stimulated *Tnf* gene expression from 72.2-fold to 23.6 fold over the unstimulated control (i.e., a 3.1-fold decrease; *p* = 0.06) in BMM from WT mice, while a non-significant increase over the unstimulated control was observed in BMM from NRF2^-/-^ mice (57.5-fold and 77.3-fold, respectively, without and with I3B). Similarly, while a 23-fold increase in *Hmox1* expression over unstimulated control was observed with WT BMM, while only a 5.2-fold increase was observed with NRF2^-/-^ BMM (i.e., a 4.4-fold decrease) (**Fig. 6A**). These findings support the involvement of NRF2 in I3B-mediated signaling in murine macrophages.

**Fig. 6.**
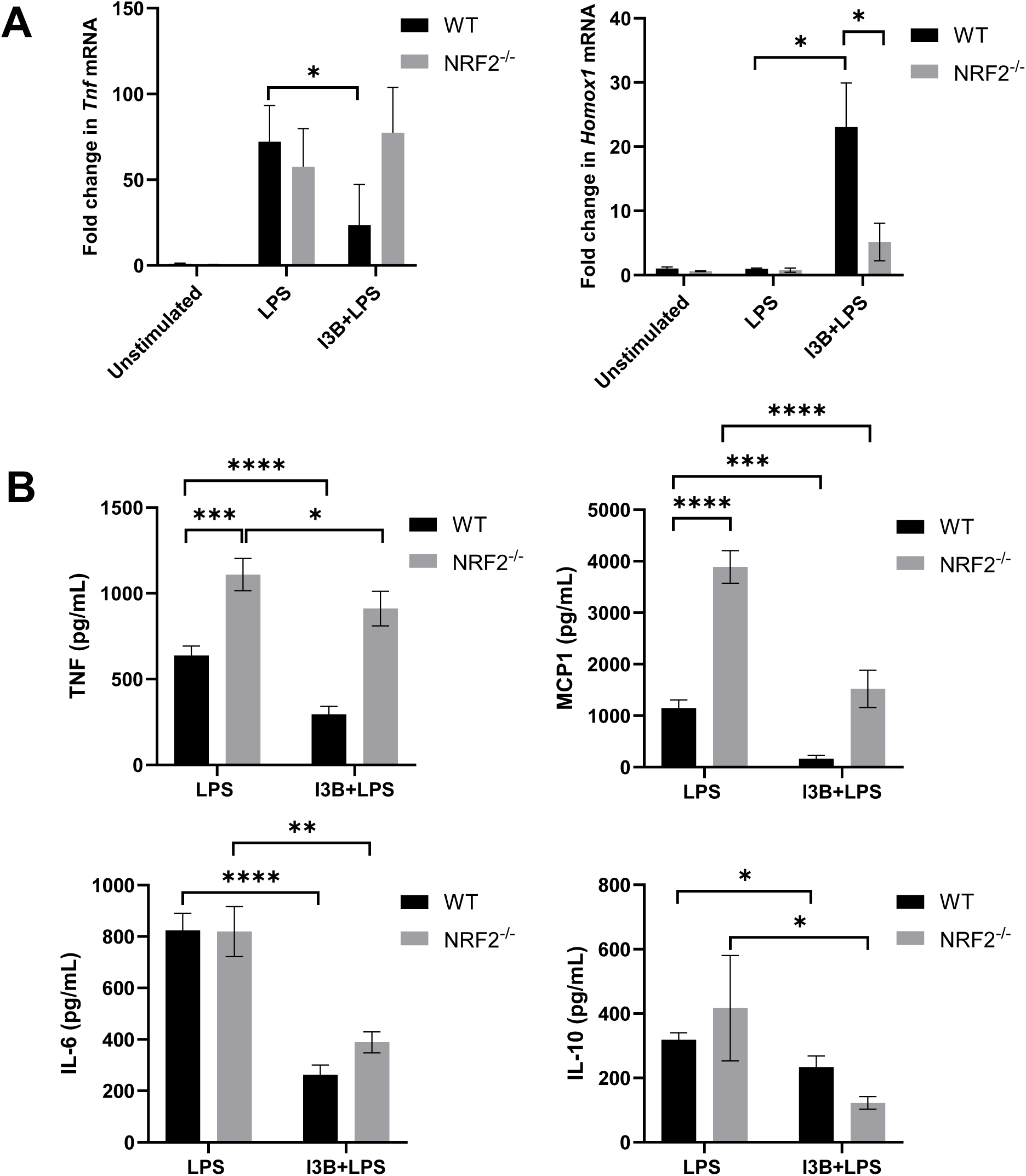
Anti-inflammatory effects of I3B in BMM are partially dependent on NRF2. (**A**) BMM from WT and NRF2^-/-^ mice were pre-incubated with 20 μM I3B for 4 h followed by stimulation with 20 ng/mL LPS for 1 h before RNA isolation. (**B**) BMM from WT and NRF2-/-were pre-incubated with 20 μM I3B, then co-stimulated with LPS for 24 h. The concentration of cytokines in culture supernatants were quantified with a multiplexed antibody bead-based assay. Values are normalized to the LPS stimulated control and are representative of three or four independent cell cultures. * *p* < 0.05, ** *p* < 0.01, *** *p* < 0.001, **** *p* < 0.0001.

Based on the observed transcriptional changes of *Tnf* and *Hmox1*, we next validated these findings by quantifying protein levels in the cell supernatant of WT and NRF2^-/-^ BMM pre-treated with I3B for 4 h followed by 24 h of 20 ng/mL LPS stimulation. The basal secretion of all the detected cytokines using the LEGENDPlex^TM^ Mouse Inflammation panel, except for IL-6, was higher in NRF2^-/-^ BMM relative to WT, suggesting a hyper-responsive phenotype in the BMM from NRF2^-/-^ mice. When normalized to the LPS control, the reduction in secreted TNF-α, MCP-1, and IL-6 with I3B pre-exposure was significantly greater in WT BMM than in NRF2^-/-^ BMM (**Fig. 6B**). On the other hand, the expression of IL-10 in WT BMM pre-incubated with 20 μM I3B was higher than NRF2^-/-^ BMM. Together, these results strongly support the hypothesis that the anti-inflammatory effects of I3B in macrophages are partially dependent on NRF2.

### I3B Modulates TNF-α Expression Independently of Gi-Protein-Coupled GPCRs

Based on our prior observations that I3B exhibits minimal cellular uptake after four hours, and studies reporting MTDMs such as kynurenic acid and tryptamine can engage GPCRs, we next examined whether this class of receptors contributes to I3B-mediated signaling. We investigated the role of Gs, Gi/αi, and Gq G-protein-coupled receptors in I3B signaling in RAW 264.7 macrophages. First, we investigated the role of Gi-coupled GPCR signaling by treating murine macrophages with pertussis toxin (PTX), a well-characterized inhibitor of Gi/αi-coupled GPCR signaling (*18*). RAW 264.7 macrophages received 200 ng/mL of PTX for 16 h, followed by treatment with 50 uM of I3B for 4 h and 100 ng/mL of LPS stimulation for 1 h. Flow cytometry analysis validated that LPS stimulation markedly increased intracellular TNF-α levels, with approximately 76.5% of macrophages positive (**Fig. 7A**). Pretreatment of cells with PTX followed by LPS (no I3B treatment) did not significantly change the proportion of TNF-α positive cells (73.8%) (**Fig. 7 A, B**). On the other hand, a comparable reduction in the proportion of TNF-α positive cells was observed when murine macrophages were treated with either I3B alone or I3B+PTX (**Fig. 7 A, B**). These results suggests that I3B suppresses LPS-induced expression of TNF-α, independent of Gi-coupled GPCR signaling.

**Fig. 7.**
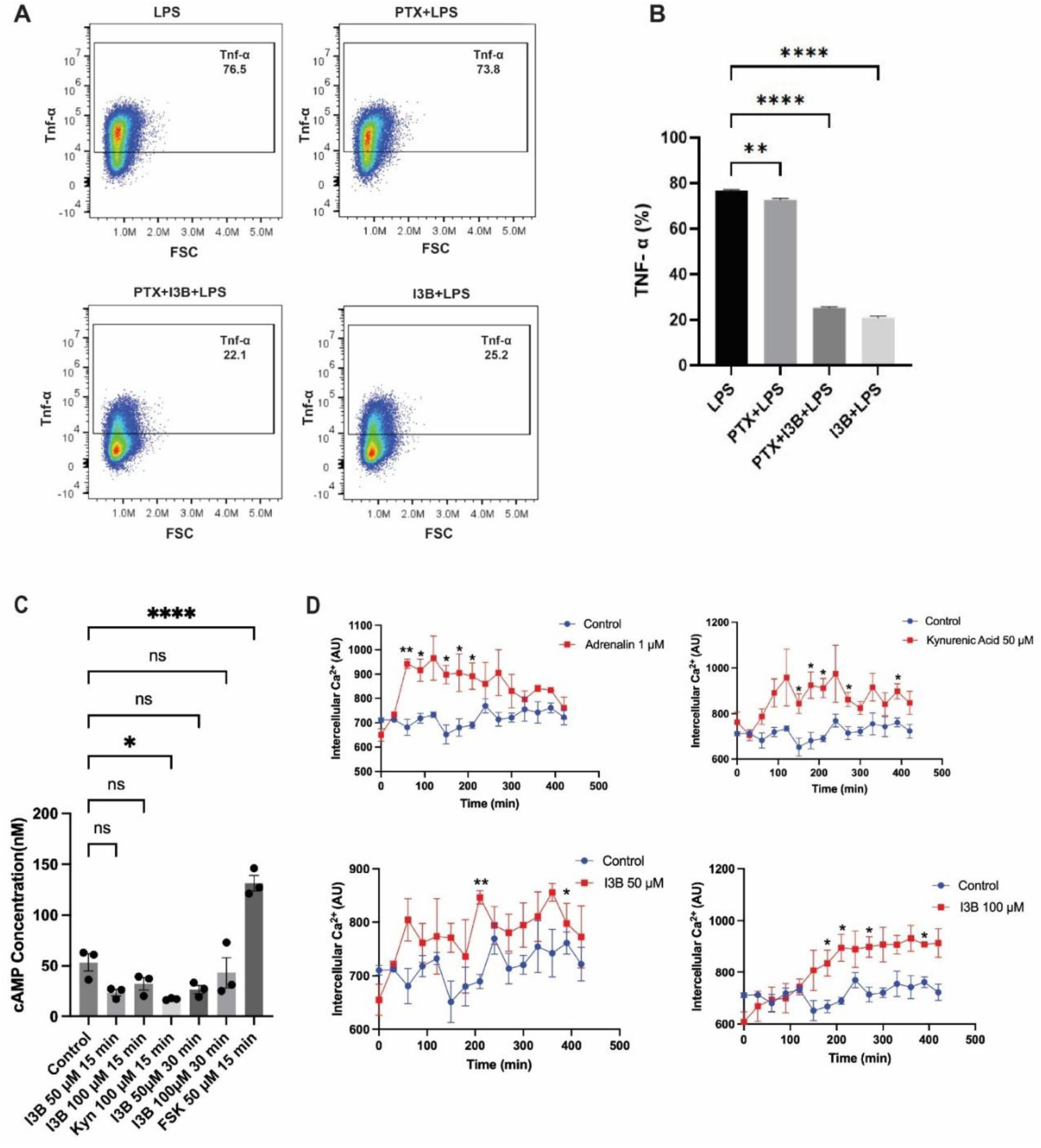
I3B attenuates inflammation dependent of Gq-coupled GPCR signaling in RAW 264.7 Macrophages. Cells were pre-treated with pertussis toxin (PTX, 200 ng/mL, 16 h) and/or I3B (50 µM, 4 h) and LPS (100 ng/mL, 1 h). (A) Intracellular TNF-α expression was assessed by flow cytometry and cytokine staining. Representative dot plots show percentage of TNF-α-positive cells indicated in each plot. (B) Quantification of TNF-α expression (mean ± SEM) from three independent experiments (n = 3). Statistical analysis was performed using one-way ANOVA followed by Tukey’s post hoc test. \*\**p* < 0.01, \*\*\*\**p* < 0.0001 indicate statistical significance compared to the LPS-treated group. (C) Intracellular cAMP levels in macrophages treated with I3B at the indicated concentrations and times compared with vehicle control and forskolin (FSK; positive control) Quantification of cAMP (mean ± SEM) from three independent experiments (n = 3). Statistical analysis was performed using one-way ANOVA \*\**p* < 0.01, \*\*\*\**p* < 0.0001 indicate statistical significance compared to the control. (D) Time-course of intracellular Ca²⁺ levels in macrophages treated with adrenaline (1 μM), kynurenic acid (50 μM), I3B (50 μM or 100 μM), compared with untreated control. Mean ± SEM from three independent experiments are shown. Statistical analysis was performed using unpaired t test. *p<0.05 \*\**p* < 0.01 indicate statistical significance compared to the control at each timepoint.

We next examined whether I3B mediates its effects through Gs- or Gi/αi-coupled receptors. Activation of Gs-coupled GPCRs stimulates adenylyl cyclase and increases intracellular cAMP, whereas Gi-coupled GPCRs inhibit adenylyl cyclase and reduce cAMP levels (*19*). To determine whether I3B engages these signaling pathways, we measured intracellular cAMP levels in LPS-stimulated cells pre-treated or not with I3B treatment. RAW 264.7 macrophages treated either 50 µM or 100 µM of I3B did not exhibit significant changes in cAMP levels compared to vehicle control (**Fig. 7C**). These findings suggest I3B does not activate GPCR signaling mediated by Gs or Gi/αi proteins. To further characterize the receptor-signaling pathway engaged by I3B, we investigated the role of Gq-coupled signaling. Activation of Gq-coupled GPCRs stimulates phospholipase C-β, leading to IP3-mediated calcium release from the Endoplasmic reticulum (*20*). Therefore, we performed a kinetic assay were we measured intracellular calcium mobilization in raw macrophages wherein we observed a significant increase in the levels of intracellular Ca²⁺ over time compared to control when treated with 100 µM of I3B exhibited (**Fig. 7D).**

The rise in intracellular Ca²⁺ together with DAG is known to activate novel PKC isoforms, including PKCδ (*21*). We therefore examined PKCδ activation by measuring the phosphorylation at Thr505, a site within the activation loop that is phosphorylated by PDK1 and serves as a marker of PKCδ activation (*22*). Treatment of RAW 264.7 macrophages with 50 µM of I3B led to a ∼2-fold increase in phospho-PKCδ (Thr505) after 5 min, which then declined in a time-dependent manner **(Fig. 8A)**. Consistent with PKCδ-dependent activation of NRF2, we found that NRF2 phosphorylation increased by ∼2-fold at 20 min after I3B (50 µM) treatment and then decreased over time **(Fig. 8B)**. Taken together, these findings support a model in which I3B signals through a Gq-coupled GPCR, leading to increased intracellular Ca²⁺, activation of PKCδ, and subsequent NRF2 phosphorylation and activation.

**Fig. 8.**
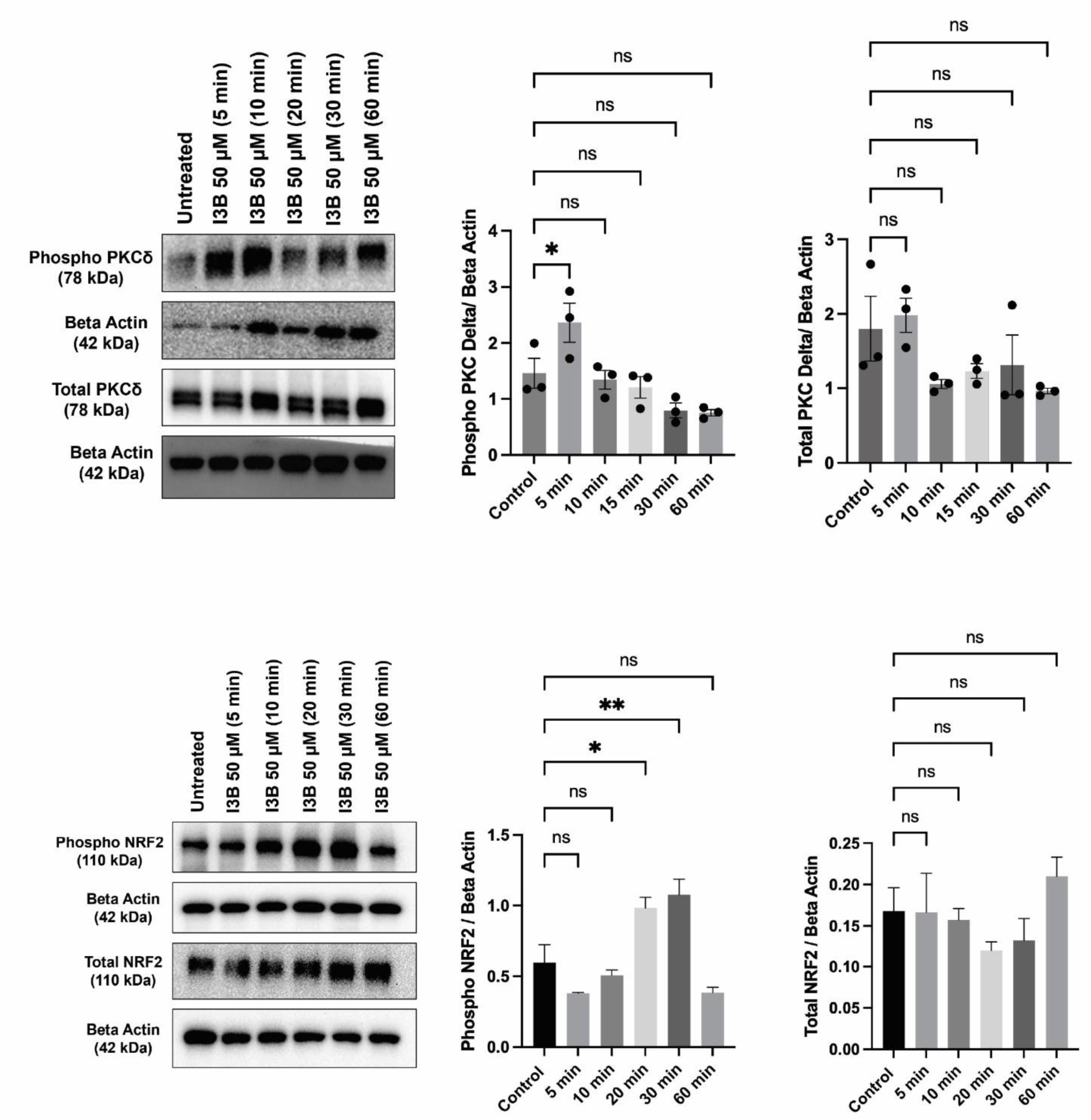
I3B treatment induces rapid PKCδ and delayed NRF2 phosphorylation in Macrophages. Cells were treated with 50 μM I3B for the indicated time points and protein expression was analyzed by Western blot. (**A**) Representative Immunoblots and densitometric quantification of phosphorylated PKCδ (p-PKCδ) and total PKCδ expression, with β-actin as loading control. p-PKCδ levels peak at 5 minutes post-treatment while total PKCδ remains unchanged. (**B**) Representative Immunoblots and densitometric quantification of phosphorylated NRF2 (p-NRF2) and total NRF2 expression. p-NRF2 levels show significant increases at 20 and 30 minutes, while total NRF2 remains stable across all time points. Data are presented as mean ± SEM (n = 3). Statistical significance was determined by one-way ANOVA with Dunnett’s post-hoc test comparing each time point to untreated control. *p < 0.05, \*\**p* < 0.01, ns = not significant.

**Fig. 9.**
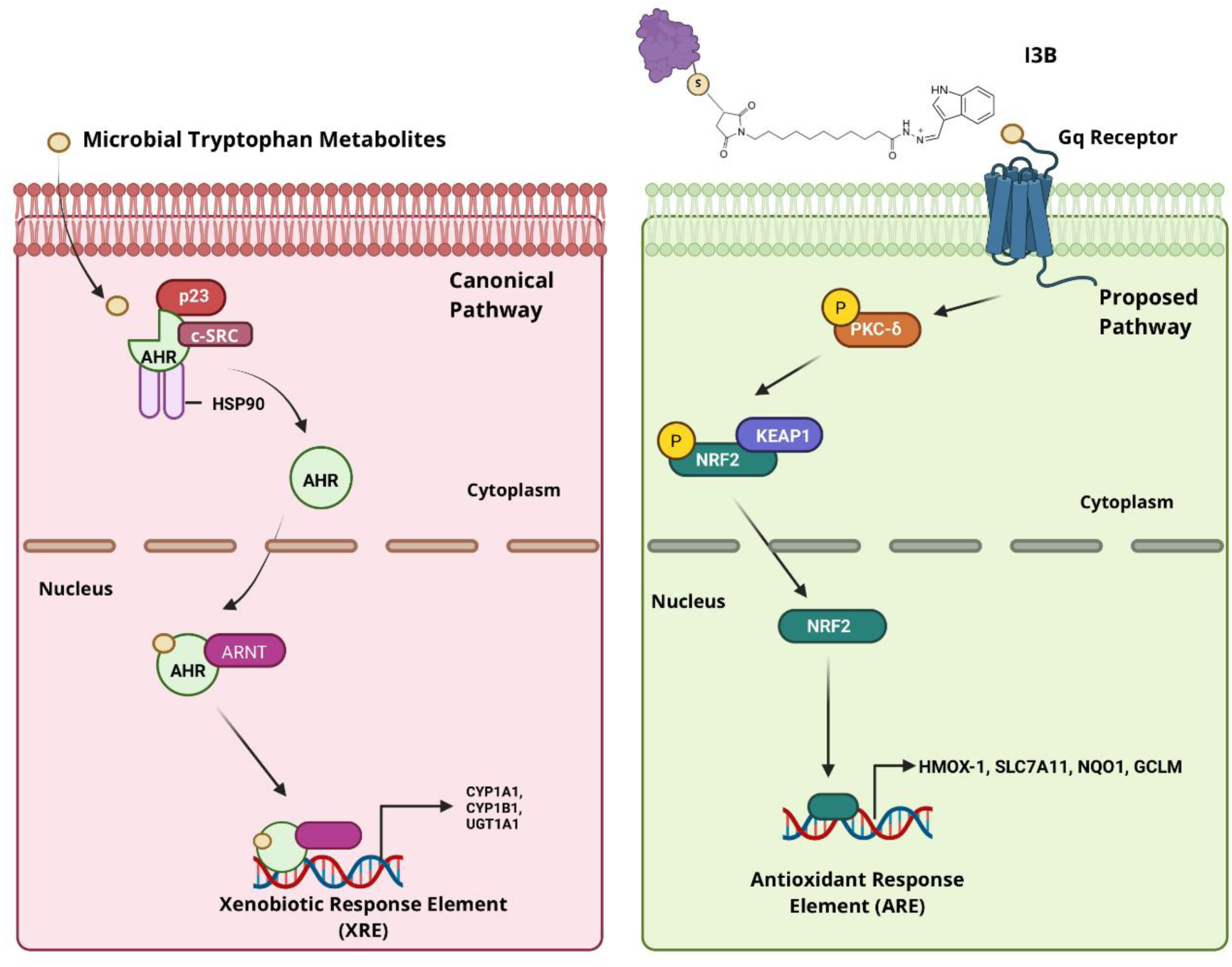
Schematic of proposed mechanisms in I3B signaling in Macrophages. Microbial tryptophan metabolites primarily translocate to the cytosol and interact directly with cytosolic receptors such as the AhR. These metabolites I3B may also engage cell surface receptor(s) but likely with a much lower frequency. On the other hand, the primary interaction of I3B is with the cell surface receptor as its intracellular is sterically hindered. This cell surface receptor-mediated signaling results in NRF2 activation and expression of downsstream genes.

## Discussion

The contribution of MDTMs to host immune function is being increasingly recognized, with evidence of several indole and indole-like compounds having pleiotropic effects on various immune cells such as epithelial and T cells (*23, 24*). Although MDTMs have been implicated in modulating inflammation, the mechanisms underlying these effects remain poorly characterized. In this study, we report that indole at physiologically relevant concentrations exerts anti-inflammatory effects in RAW 264.7 macrophages. Furthermore, using a cell-impermeant indole conjugate, we provide evidence that suggest these effects can be mediated by an as-yet unidentified cell surface receptor.

Many prior studies investigating the mechanism of MDTMs have focused on AhR-dependent responses. Indole acts as a weak AhR agonist and an antagonist of 2,3,7,8-tetrachlorodibenzo-p-dioxin (TCDD)-stimulated AhR responses in epithelial cells and hepatocytes (*25*). Some of the protective effects of IAld on colonic inflammation have been shown to be AhR dependent (*26, 27*). However, the comparable reduction in TNF-alpha levels in WT and AhR^-/-^BMM following indole treatment, together with the inability of an AhR antagonist to affect indole-mediated TNF-α suppression in RAW264.7 cells, suggests that indole acts through an AhR-independent mechanism in both macrophage models. Supporting this, previous studies have also reported that AhR expression is markedly lower in RAW264.7 macrophages compared to murine AML12 hepatocytes (*10, 28*). Collectively, these findings highlight the significance of cell type–specific signaling responses to MDTMs, emphasizing the need to consider cellular context when evaluating their mechanisms of action.

To further probe MTDMs AhR-independent effects we synthesized I3B, a cell-membrane impermeant analog of indole through conjugation of IAld to BSA. Interestingly, I3B was approximately 50-fold more potent compared to indole, in attenuating LPS-mediated induction of *TNF-α* in RAW264.7 macrophages. I3B’s attenuation of LPS-induced cytokine production in BMM extend beyond just *TNF-α*, *MCP-1* and *IL-6* levels were also reduced. These findings suggest that I3B exerts its anti-inflammatory effects through engagement of cell surface receptors rather than intracellular targets, implicating a membrane-initiated signaling mechanism in the attenuation of LPS-induced inflammatory responses. Furthermore, the relative reduction in secretion of these pro-inflammatory cytokines was more pronounced in WT compared to NRF2^-/-^ BMM. Suggesting the anti-inflammatory potential of I3B is partially dependent on NRF2. To further support this, the relative levels of secreted anti-inflammatory *IL-10* were higher in the supernatants of WT compared to NRF2^-/-^ BMM.

NRF2 is a transcription factor that regulates cellular responses to numerous stimuli, including oxidative and electrophilic stress, heme metabolism, detoxification of xenobiotic compounds, and inflammation (*29, 30*). Under basal conditions, NRF2 is sequestered in the cytoplasm by its inhibitor, Kelch-like ECH-associated protein 1 (Keap1). Activation of NRF2 results in its dissociation from Keap1 and allows for nuclear translocation and up-regulation of ARE-regulated genes (*31, 32*). Transcriptome profiling of I3B treated LPS-stimulated RAW264.7 macrophages identified the upregulation of ARE-regulated phase II enzyme genes: Hmox1, *Slc7a11*, *Nqo1*, and *Gclm*. These genes encode enzymes which functions include to modify and eliminate numerous exogenous and endogenous compounds (*33*). Beyond eliminating ROS, NRF2 modulates inflammation via independent pathways (*34, 35*). Furthermore, direct binding of NRF2 to non-ARE sites in the proximity of LPS-regulated cytokines such as IL-6 and *IL1-β* has been shown to inhibit their expression and RNA polymerase recruitment in macrophages (*36*).

Of the ARE-regulated genes upregulated by I3B, Hmox1 demonstrated the highest induction (18-fold). The primary enzymatic function of heme-oxygenase 1 (*HO-1*), the enzyme encoded by Hmox1, is to catalyze the conversion of free heme to biliverdin, releasing carbon monoxide and ionic iron as by-products (*37*). In addition to detoxifying accumulating heme, HO-1 contributes to anti-inflammatory signaling via the release of endogenous carbon monoxide (22), possibly through an IL-10 related mechanism (*38*). The potent induction of Hmox1 led us to hypothesize that the anti-inflammatory effects of I3B may be mediated directly by HO-1 activity. However, experiments with a competitive inhibitor of *HO-1*, tin protoporphyrin IX, did not result in significant changes in I3B-mediated reduction of LPS-stimulated TNF-α accumulation (data not shown). A similar lack of effect was observed using an IL-10 neutralizing antibody (data not shown), suggesting that induction of HO-1 alone may not be sufficient to account for the anti-inflammatory effects observed with I3B.

Since NRF2 is a transcription factor that is localized in the cytoplasm when inactive (*39*), direct engagement of NRF2 with the cell membrane impermeant I3B is unlikely. This supports the involvement of a novel surface-mediated signaling mechanism in mediating the anti-inflammatory effects of I3B in macrophages. The observation that I3B inhibits LPS-stimulated TNF accumulation at concentrations 50-fold lower than free indole suggests a model where cell surface engagement of indole triggers more potent anti-inflammatory signaling than what free indole does through engagement with intracellular receptors. When cells are exposed to indole, a kinetic balance exists between indole binding to cell-surface receptors and intracellular uptake via diffusion or active transport. In this scenario, rapid intracellular transport likely limits indole’s interaction with cell surface receptors. However, when cells are exposed to I3B, the molecule’s larger size impedes intracellular transport, which shifts the balance toward interaction with cell-surface receptors, resulting in sustained receptor engagement and the observed potent effects.

GPCRs constitute the largest receptor superfamily in humans, comprising over 800 members and playing crucial roles in diverse physiological and pathological processes (*40*). Transcriptional profiling analyses have confirmed that macrophages express a variety of GPCRs and immune cells, such as leukocytes, express numerous GPCRs that are involved in essential biological functions, including chemotaxis, cell migration, and inflammatory responses (*39, 41*). The tryptophan-derived metabolite tryptamine has been shown to interact with multiple GPCRs, including serotonin receptors (5-HT1A, 5-HT2A, and 5-HT2C), trace amine-associated receptor 1 (TAAR1), and GPRC5A influencing neurological and immune responses (*42, 43*). Kynurenine-derived metabolites, notably kynurenic acid, have also been reported as endogenous ligands for G protein-coupled receptor 35 (*14, 44*) that primarily couples to Gαi/o and Gα_12/13_ proteins to regulate inflammatory and nociceptive signaling pathways (*45*). However, our findings suggest that Gα_i_ receptors are not engaged by I3B, suggesting the involvement of other GPCRs in mediating I3B signaling in RAW264.7 macrophages. Another potential class that could mediate I3B’s signaling cascade is Gαs-coupled GPCRs. The primary downstream effector of Gs receptors is adenylyl cyclase, an enzyme that catalyzes the conversion of adenosine triphosphate (ATP) to cyclic adenosine monophosphate (cAMP) (*46*). Since blocking Gs protein signaling before I3B treatment of murine macrophages did not significantly alter levels of cAMP, our results suggest that I3B effects are not mediated through a Gs coupled receptor. This allowed us to further probe the mechanism of action of another group of G protein coupled receptors, Gq class of GPCRs.

Gq proteins activate phospholipase C-β (PLC-β), which hydrolyzes phosphatidylinositol 4,5-bisphosphate (PIP₂) to generate the second messengers diacylglycerol (DAG) and inositol 1,4,5-trisphosphate (IP₃), leading to intracellular calcium release and protein kinase C (PKC) activation (*47*). The significant increase in cytosolic calcium levels compared to control suggest that I3B mediates its effects through a Gq-coupled signaling pathway. This was further confirmed, when we also observed an increase in PKCδ phosphorylation, a downstream effector of Gq protein signaling, in RAW 264.7 macrophages following I3B treatment. Gq signaling activates phospholipase C-β, which leads to IP3 and diacylglycerol (DAG) generation and increased intracellular Ca²⁺ [35]. The rise in intracellular Ca²⁺ coupled with DAG generation is known to activate novel PKC isoforms, including PKCδ which is activated by phosphorylation at Thr505. Importantly, activated PKC isoforms can directly phosphorylate NRF2 at Ser40, disrupting its interaction with Keap1 and facilitating its nuclear translocation and transcriptional activity (*48*).

PKCδ has been identified as a direct NRF2 kinase that phosphorylates NRF2 at serine 40 (Ser40) (*49, 50*). This phosphorylation event is a critical step for NRF2 activation in response to antioxidant compounds and oxidative stress. I3B’s signaling through a Gq-coupled GPCR protein leading to PKCδ activation and subsequent NRF2-mediated responses suggest a sophisticated cellular defense mechanism linking extracellular stimuli to coordinated antioxidant and anti-inflammatory responses. In mammals, there are four Gq-family α-subunits(*51*). Therefore, further studies are needed to identify which Gq GPCR is involved in mediating I3B signaling that leads to NRF2 activation.

Our confocal microscopy data show localization of indole conjugated to NeutrAvidin at discrete regions on the cell surface, rather than being distributing uniformly across the membrane. This spatially restricted localization is consistent with well-characterized mechanisms of GPCR organization. GPCRs and associated signaling molecules preferentially partition into lipid rafts, cholesterol, and sphingolipid-enriched membrane micro domains. These domains serve as platforms for the assembly of signaling complexes and bring receptors into proximity with heterotrimeric G proteins, kinases, and other effector molecules (*52*). Such compartmentalization has been documented for multiple GPCR families, including chemokine receptors and metabolite-sensing receptors, which preferentially cluster in these domains(*53, 54*). These organized structures enhance signaling efficiency and fidelity by increasing the local concentration of receptors and their transducers, allowing cells to respond in a coordinated manner to specific ligands (*55*).

The crosstalk between NRF2 and nuclear factor kappa B (NF-κB) signaling pathways suggests potential mechanisms for describing the observed of antioxidant and anti-inflammatory responses. NRF2 negatively regulates the NF-κB signaling pathway, a master regulator of inflammatory gene expression, through multiple mechanisms (*56*). First, NRF2 inhibits oxidative stress-mediated NF-κB activation by decreasing intracellular ROS levels through upregulation of antioxidant enzymes (*50*). Second, NRF2 prevents proteasomal degradation of IκB-α, the cytoplasmic inhibitor of NF-κB, thereby blocking nuclear translocation of NF-κB (*57*). Emerging evidence also shows that NRF2 can also directly suppress transcription of pro-inflammatory cytokine genes, representing a mechanism independent of its antioxidant function. Studies also demonstrate that NRF2 interferes with lipopolysaccharide (LPS)-induced transcriptional upregulation of pro-inflammatory cytokines including interleukin-6 (IL-6) and interleukin-1β (IL-1β) (*58, 59*) through disruption of RNA polymerase II (Pol II) recruitment to pro-inflammatory gene loci (*60*). This transcriptional interference is independent of ROS levels and antioxidant response elements, indicating a distinct anti-inflammatory function of NRF2 separate from its antioxidant activities (*61*).

In summary, we report a novel our results demonstrate that I3B exerts anti-inflammatory effects in murine macrophages through Gq-coupled GPCR signaling and intracellular Ca^2+^ mobilization, independent of Gi-coupled GPCRs and the AhR. Instead, I3B’s anti-inflammatory effects are partially dependent on NRF2. Future work includes employing receptor-specific antagonists or genetic knockout models targeting NRF2 and alternative GPCR pathways, to elucidate the underlying molecular mechanisms. Identification of cell surface receptor(s) that are involved in mediating the effects of indole and other MDTMs could also potentially lead to new strategies for using MDTMs or their synthetic derivatives as therapeutic molecules.

## Materials and Methods

### Animals and cell culture

RAW264.7 macrophages were purchased from ATCC (Manassas, VA) and cultured in DMEM supplemented with glucose, penicillin, streptomycin, non-essential amino acids, HEPES, and 10% FBS at 37 °C and 5% CO_2_ on 10 mM tissue-culture treated dishes. Cells were trypsinized for passage at roughly 80% confluence and seeded into tissue culture plates for experiments. Cells seeded in well-plates were cultured overnight before exposure to experimental conditions.

For AhR experiments, C57B/6 mice-derived wild-type and AhR^-/-^ mice (approximately 6-8 weeks of age) were purchased from Taconic. All experiments were carried out after 2 weeks of acclimatization in the LARR animal facility at Texas A&M (AUP #2017-0145). For NRF2^-/-^experiments, C57B/6 mice-derived wild-type and NRF2^-/-^ mice femurs and tibias were provided by Dr. Venkatakrishna Rao Jala (University of Louisville). BMM were cultured as previously described (*62*). Briefly, cells were flushed from the femurs and tibias of mice and cultured in 10 mM non-tissue culture treated dishes with RPMI containing 10 ng/mL macrophage colony-stimulating factor at 37 °C and 5% CO_2._ Cells were fed additional media after three days of culture and harvested after seven days.

### Flow cytometry

RAW264.7 cells were seeded at a density of ∼2.7 −3.2 x 10^4^ cells per well in a round bottom TC-treated 96 well plate. Culture media was removed, and cells were washed with 200 μL cold PBS supplemented with 5% BSA twice and fixed with 200 μL 4% PFA for 20 minutes on ice. Cells were permeabilized with 200 μL BD Perm/Wash Buffer (BD Biosciences) for 20 minutes. Non-specific antigen binding was blocked with 0.1 μg Anti-Mouse CD16/CD32 (BD Biosciences, CA) in 75 μL for 5 minutes before cells were stained with TNF Rat anti-Mouse, Alexa Fluor 700, Clone: MP6-XT22 (BD Biosciences) for 30 minutes. Stained cells were resuspended in PBS and analyzed in a BD LSRFortessa™ X-20 and processed in FlowJo software (BD Biosciences).

To evaluate Gαi involvement in I3B signaling, RAW 264.7 cells were pre-treated with pertussis toxin (PTX, 200 ng/mL) for 16 hours, followed by I3B (50 µM, 4 hours) and LPS (100 ng/mL, 1 hour). After stimulation, cells were stained with LIVE/DEAD™ Fixable Aqua (Thermo Fisher, L34957) for 30 minutes at room temperature to exclude non-viable cells, followed by Fc receptor blocking using BD Fc Block™ (CD16/32, BD Biosciences, 553141) for 10 minutes at 4°C. Surface staining was performed in FACS buffer (1× PBS + 2% FBS) with fluorophore-conjugated antibodies for 30 minutes at 4°C. Cells were then fixed with 2% paraformaldehyde (Thermo Fisher, J19943.K2) for 20 minutes and permeabilized using BD Perm/Wash Buffer (BD Biosciences, 554723) for 20 minutes at 4°C. Intracellular TNF-α (AF-410-NA, R&D Systems) was stained using fluorochrome-conjugated antibodies in Perm/Wash buffer and incubated for 30 minutes. Data were acquired on MoFlo Astrios EQ (Beckman Coulter), BD LSRFortessa™ X-20 (BD Biosciences), or Cytek® Aurora (Cytek Biosciences). Gating and compensation were performed using fluorescence minus one (FMO) and unstained controls. Data analysis was conducted using FlowJo Version 10 software.

### Synthesis of I3B

Indole-3-aldehyde (IAld) dissolved in dimethyl formamide (DMF) was reacted with KMUH in DMSO at a 5:1 molar ratio at room temperature on a shaker for 2 h. BSA was dissolved in PBS supplemented with 2 mM EDTA and the pH adjusted to 6.8. The concentration of this solution was used to calibrate BSA protein quantification using a Nanodrop 2000 by measuring absorbance at 280 nm wavelength. The KMUH-IAld reaction product was reacted with BSA at a 5:1 molar ratio (KMUH equivalents to BSA) on a shaker at room temperature for 2 h. Unreacted reagents were removed with a ZEBA desalting column (ThermoFisher) prepared according to manufacturer recommendations. The desalted product was concentrated in an Amicon® Ultra filter unit (3 kDa cutoff) prepared according to manufacturer recommendations at 4300 x *g* in a swinging-bucket rotor centrifuge for 1 h. The volume of the concentrated product quantified using a Nanodrop 2000. Buffer-exchange with serum-free DMEM was carried out in the Amicon® Ultra at 4300 x *g* column in a swinging-bucket rotor centrifuge for 1 h. The volume of the concentrated product was measured, and fetal bovine serum (FBS) added to final concentration of 10%. The I3B concentrate was then sterilized through a polyethersulfone 0.45 µM syringe filter. Desalted I3B without buffer exchange was used for MALDI mass spectrometry analysis. Samples were analyzed in an AXIMA MALDI-TOF Mass Spectrometer (Shimadzu) in reflectron mode at the Protein Chemistry Laboratory, Texas A&M University.

### Synthesis and Cellular Imaging of Indole-NeutrAvidin Conjugate

Indole-3-aldehyde (IAld) was first dissolved in DMSO and reacted with 4-aminobenzoic acid hydrazide at a 5:1 molar ratio under gentle agitation at room temperature for 4 hours. Subsequently, biotin-NHS was added to the reaction mixture at a 10:1 molar ratio (relative to hydrazide) and incubated under similar conditions for an additional 4 hours. NeutrAvidin-FITC (Thermo Fisher Scientific) was reconstituted in PBS to a concentration of 10 mg/mL and added to the biotinylated IAld mixture at a 10:1 molar ratio. The reaction was carried out overnight at 4 °C on a shaker to facilitate conjugation. Unreacted small molecules were removed using a Zeba™ desalting column (Thermo Fisher Scientific), prepared and used according to the manufacturer’s protocol. The conjugated product was then concentrated using an Amicon® Ultra centrifugal filter unit (3 kDa MWCO) at 4,300 × *g* for 1 hour at 4 °C in a swinging-bucket rotor. The volume of the concentrated conjugate was quantified using a NanoDrop™ 2000 spectrophotometer. Buffer exchange into serum-free DMEM was subsequently performed using the same filter unit under identical centrifugation conditions. The final volume was measured, and fetal bovine serum (FBS) was added to a final concentration of 10%.

For cellular imaging, approximately 2 × 10^⁶^ cells in 2 mL of medium were seeded into 30 mm glass-bottom dishes and incubated with 20 µM of the fluorescent Indole-NeutrAvidin-FITC conjugate. After incubation, the cells were washed twice with PBS and stained with CellMask™ Deep Red Plasma Membrane Stain (Thermo Fisher Scientific) following the manufacturer’s protocol. The stained cells were washed again with PBS, replaced with ultrapure water, and imaged using a Zeiss LSM 880 confocal microscope equipped with Airyscan (Zeiss, White Plains, NY). Fluorescence imaging was conducted using excitation/emission settings of 649/666 nm for membrane visualization (red channel) and 494/518 nm for the Indole-NeutrAvidin-FITC conjugate (green channel). Image acquisition and analysis were performed using Zeiss image analysis software.

### Multiplexed cytokine analysis

RAW264.7 cells were cultured as previously stated for ICS-FACS. Supernatants were collected at the end of 24 h LPS stimulation and centrifuged at 4 °C, 400 RCF for 10 minutes. Supernatants were separated from any debris and stored at −80 °C before analysis. Samples were assayed with a LEGENDplex™ Mouse Inflammation Panel (13-plex) with the use of a filter plate according to manufacturer recommendations. Samples from the bead-based assay were then analyzed with a BD LSRFortessa™ X-20 and data was processed using BioLegend’s LEGENDplex Data Analysis Software.

### Quantitative Reverse Transcription PCR

Cultured RAW 264.7 cells were rinsed with cold PBS and frozen at −80 °C before RNA isolation with E.Z.N.A. Total RNA Kit I, (Omega Bio-tek). Cells were lysed according to manufacturer protocols and homogenized with Omega Homogenizer Columns. Total RNA was quantified in a Nanodrop 2000. RT-qPCR was performed using qScript One-Step SYBR Green reagents in a Light cycler 96 System.

### RNA Sequencing

Total were extracted from RAW264.7 cells (n = 4) using E.Z.N.A. ® Total RNA Kit I. RNA integrity was assessed using Agilent 2200 TapeStation. Samples were prepared using TrueSeqRNA sample preparation for paired-end 150 bp reads in an Illumina NextSeq 500 (Molecular Genomics Workspace, Texas A&M University, College Station, TX). Adapter sequences were removed with Trimmomatic, version 0.39 allowing no mismatches and removing leading and trailing sequences with a quality score below 9. Trimmed sequences were mapped to the Mouse genome assembly GRCm39 obtained from NCBI and indexed in STAR. Raw counts were normalized using the trimmed mean of M values and differential gene expression were analyzed using EdgeR. Gene set enrichment analysis and leading-edge analysis were performed using GSEA software from MSigDB (*63, 64*).

### Intracellular Ca²⁺ assays

Intracellular calcium was measured in macrophages using the Fluo-8 Calcium Flux Assay Kit (Abcam, ab112129) following the manufacturer’s protocol.

### cAMP assay

Cellular cAMP was quantified with the cAMP Assay Kit (Competitive ELISA, fluorometric; Abcam, ab138880). Briefly, 4E+04 macrophages were seeded per well in a 96-well plate, and all steps were performed according to the manufacturer’s instructions.

### Western blot analysis

Macrophages (∼4E+05 cells per well in a 6-well plate) were treated with the indicated compounds for the specified durations and lysed in modified RIPA buffer supplemented with protease inhibitor cocktail (Pierce), 10 mM NaF, and 1 mM Na_3_VO_4_. Protein concentration was determined by BCA assay (Pierce). Samples were denatured with SDS and ∼20 µg total protein was resolved on a 10% SDS-PAGE gel, then transferred to a PVDF membrane (Thermo Fisher Scientific, Inc.) by wet transfer. Membranes were blocked with EveryBlot Blocking Buffer (Bio-Rad) and probed with primary antibodies against phospho-PKCδ (Cell Signaling Technology9374), total PKC (Cell Signaling Technology 2508), and β-actin (Cell Signaling Technology 12620), followed by HRP-conjugated anti-rabbit secondary antibody (Cell Signaling Technology 7074). Signals were developed using Clarity Max Western ECL substrate (Bio-Rad) and imaged on a ChemiDoc system (Bio-Rad). Band intensities were quantified in ImageJ and normalized to β-actin from the same lane.

## Acknowledgments

We thank Dr. Lawrence Dangott (Protein Chemistry Laboratory, Texas A&M University) for discussions on I3B characterization. This work was partially supported by the Ray B. Nesbitt endowed chair (to AJ).

## Funding

National Institutes of Health grant 1 R01 AI110642-01 (AJ and RCA)

## Author contributions

Conceptualization: CC, SS, AJ

Methodology: AJ, CC, RCA, SRS, AM, SG, VJ

Investigation: CC, SRS, AM Supervision: AJ, RCA, SS

Writing original draft: CC, AJ, RCA, SRS

Writing review & editing: AJ, CC, RCA, SRS, AM, YSM

## Competing interests

Authors declare that they have no competing interests.

## Notes

### Competing Interest Statement

The authors have declared no competing interest.

