## Supplementary Figures and Tables for "NRF2-Dependent Anti-Inflammatory Activity of Indole via Cell Surface Receptor Signaling in Murine Macrophages"

Clint Chen et al.

### Supplementary Text

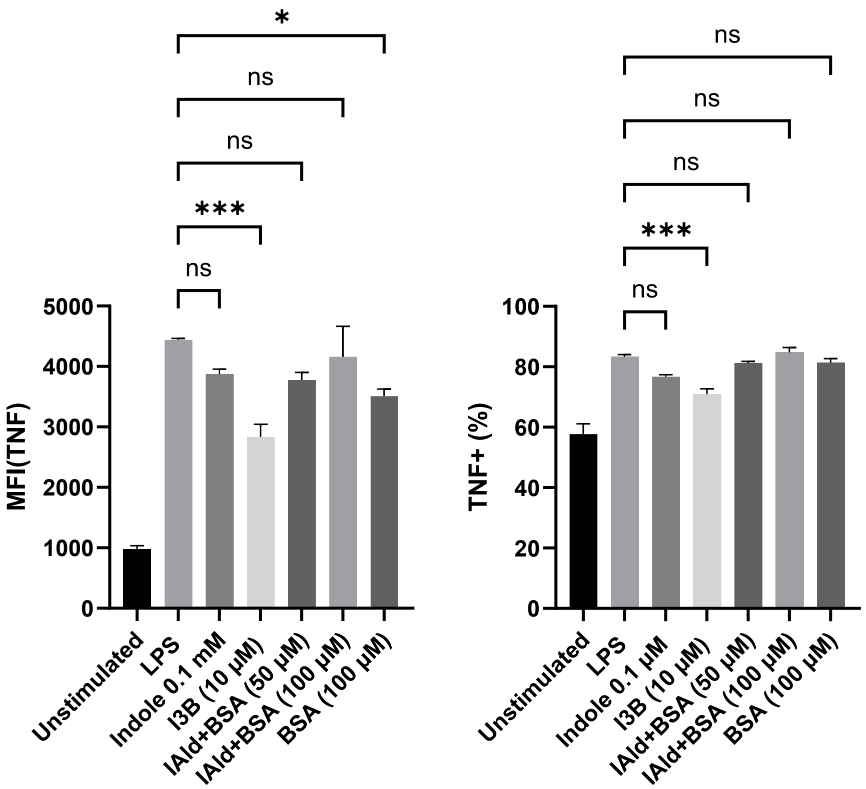


**Fig. S1. Anti-inflammatory Effects of Indole, IAld–BSA (50 and 100 µM), and BSA**
RAW264.7 macrophages were pre-incubated with compounds of interest for 4 h followed by co-stimulation with 250 ng/mL of LPS for 6 h and ICS flow cytometry. Quantification of TNF-α protein was carried out using the gating strategy shown in **Figure 1(A)**. Mean fluorescence intensity (arbitrary units) and percent TNF-α positive cells are shown. Values are normalized to the LPS stimulated control and are representative of three or four independent cell cultures. * $p < 0.05$, ** $p < 0.01$, *** $p < 0.001$, **** $p < 0.0001$.

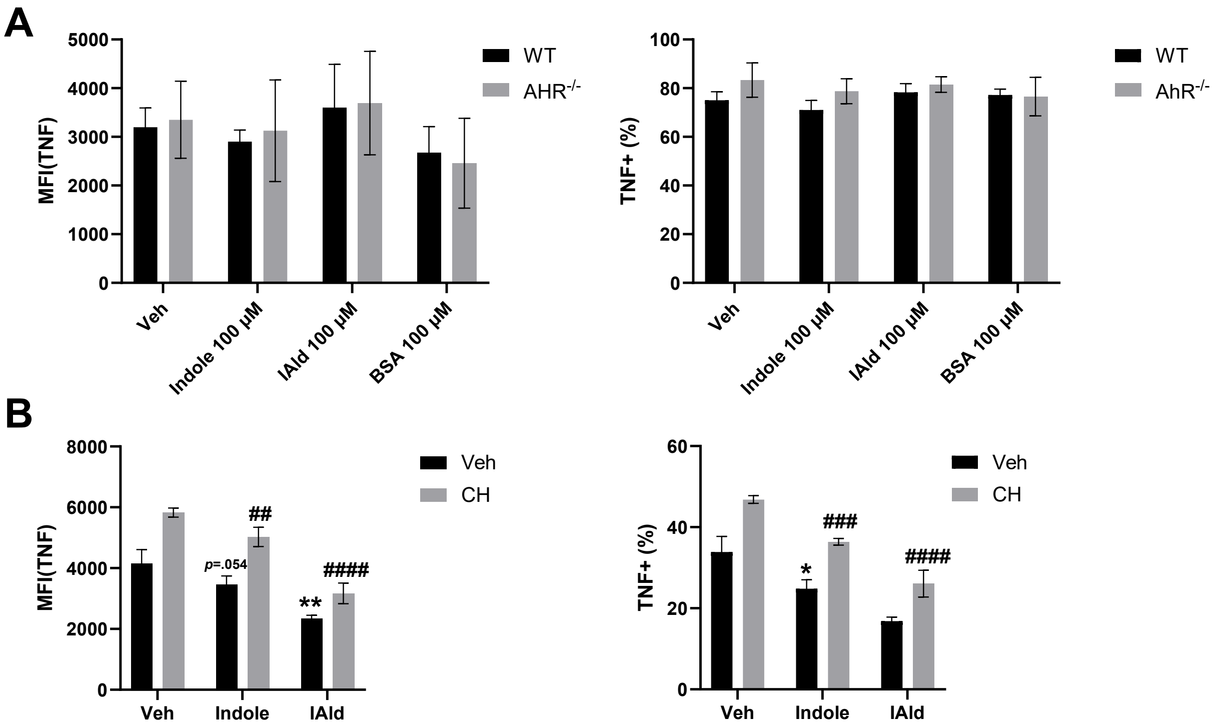


**Fig. S2. Effects of I3B on macrophages is AhR independent.**

**(A)** Bone marrow-derived macrophages (BMMs) from wild-type (WT) and AhR knockout ($AhR^{-/-}$) mice were treated with indole, IAld or controls for 4 hours, followed by LPS stimulation (20 ng/mL) for 6 hours. TNF-α levels were assessed by ICS flow cytometry. Absolute MFI (left) and percent TNF-α–positive cells (right) are shown. **(B)** RAW264.7 macrophages were pre-treated with the AhR antagonist CH-223191 (CH; 20 μM) for 14 hours, followed by co-incubation with indole, indole-3-aldehyde (IAld) (all at 50 μM), or vehicle control (Veh) for 4 hours. Cells were subsequently stimulated with LPS (250 ng/mL) for 6 hours and analyzed by intracellular cytokine staining (ICS) and flow cytometry. The absolute mean fluorescence intensity (MFI) values and percentages of TNF-α–positive cells. Data are presented as mean ± SEM. *$p < 0.05$, **$p < 0.01$, ***$p < 0.001$ relative to vehicle control (A) or WT (B); #$p < 0.05$, ##$p < 0.01$, ###$p < 0.001$ relative to vehicle + CH condition (A) or $AhR^{-/-}$ (B); ns = not significant.

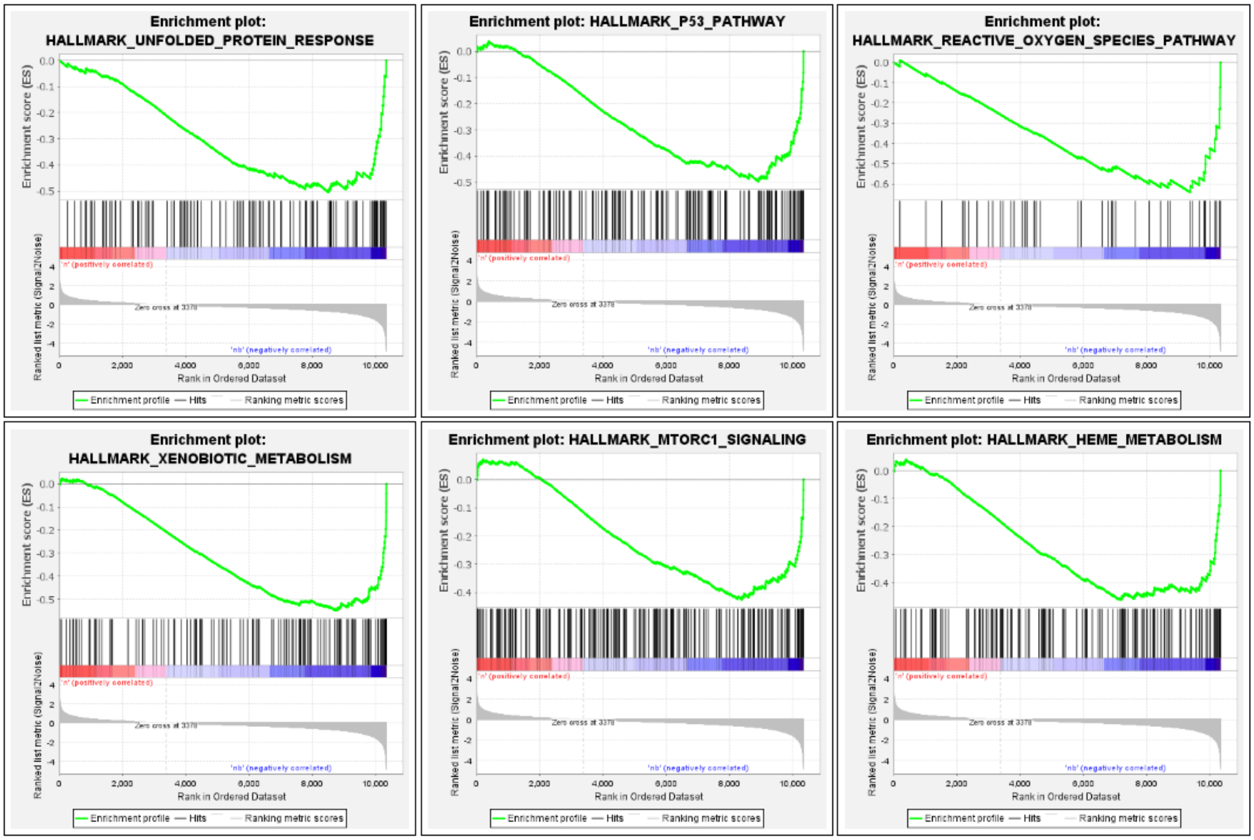


**Fig S3**. **Top gene sets enriched by I3B in Wikipathways collection.** Gene Set Enrichment Analysis (GSEA) reveals significant negative enrichment of six hallmark pathways in the treatment group:(A) Unfolded Protein Response,(B) P53 Pathway,(C) Reactive Oxygen Species Pathway,(D) Xenobiotic Metabolism, (E) mTORC1 Signaling, and (F) Heme Metabolism.

**Table S1**. **Primer sequences for qRT-PCR analysis**

| ***Gene*** | ***Forward Primer Sequence*** | ***Reverse Primer Sequence*** |
|---|---|---|
| *Hmox1* | 5'-GGTGATGGAGCGTCCACA-3' | 5'-AGGAAGCCATCACCAGCTTAA -3' |
| *Tnf-α* | 5'-TCTCATGCACCACCATCAAGGACT-3' | 5'-TGACCACTCTCCCTTTGCAGAACT-3' |
| *Cxcl10* | 5'- CCAAGTGCTGCCGTCATTT-3' | 5'- TTTCATCGTGGCAATGATCTCA-3' |

**Table S2. Top differentially regulated genes in LPS stimulated RAW264.7 cells.**

| Gene | Fold Change | p-value |
|---|---|---|
| Cd69 | 90.6 | 2.70E-18 |
| Cxcl2 | 88.6 | 7.10E-41 |
| Cxcl10 | 40.3 | 9.40E-38 |
| Csf3 | 24.7 | 1.40E-21 |
| Egr2 | 21.6 | 1.50E-31 |
| Tnfaip3 | 19.6 | 3.30E-43 |
| Ccrl2 | 19.6 | 1.30E-27 |
| Ppp1r15a | 16.8 | 1.80E-38 |
| Egr1 | 13.4 | 8.10E-42 |
| Ifnb1 | 12.3 | 2.80E-13 |
| Myc | 12.9 | 3.30E-35 |
| Egr3 | 11.6 | 3.30E-35 |
| Tnf 10.1 | 10.1 | 1.50E-41 |
| Zfp36 | 10 | 5.00E-43 |
| Socs3 | 9.7 | 1.30E-37 |
| Ccl4 | 9.4 | 1.70E-35 |
| Edn1 | 8.1 | 1.10E-12 |

**Table S3. Top differentially regulated genes in I3B treated RAW264.7 cells.**

| Gene | Fold Change | p-value |
|---|---|---|
| Sel1l3 | 34.48 | 2.80E-09 |
| Cox6a2 | 23.22 | 1.10E-08 |
| LOC118567915 | 0.05 | 1.20E-12 |
| Slc40a1 | 20.37 | 3.30E-25 |
| Hmox1 | 17.69 | 7.10E-28 |
| Abca8b | 16.92 | 1.00E-07 |
| LOC118568257 | 16.16 | 1.50E-06 |
| Cttn | 0.08 | 1.00E-08 |
| Trib3 | 12.36 | 2.40E-26 |
| Aqp9 | 11.54 | 8.70E-18 |
| Slc1a4 | 9.68 | 1.80E-28 |
| Gm40960 | 0.1 | 1.20E-11 |
| Adh7 | 9.53 | 2.70E-14 |
| Extl1 | 9.45 | 1.90E-12 |
| Aox1 | 9.37 | 2.20E-12 |
| Slc6a9 | 7.93 | 6.50E-28 |
| Slc7a11 | 7.61 | 3.00E-34 |
